# An allosteric network governs Tom70 conformational dynamics to coordinate mitochondrial protein import

**DOI:** 10.1101/2025.07.19.665690

**Authors:** Max J. Bachochin, Kelly L. McGuire, Brian D. Cook, Qiaozhen Ye, Steve Silletti, Kevin D. Corbett, Elizabeth A. Komives, Mark A. Herzik

**Affiliations:** Department of Chemistry and Biochemistry, University of California San Diego, La Jolla, CA, 92093 USA; Department of Cellular and Molecular Medicine, University of California San Diego, La Jolla, CA, 92093 USA; Department of Molecular Biology, University of California San Diego, La Jolla, CA, 92093 USA

## Abstract

Tom70 mediates mitochondrial protein import by coordinating the transfer of cytosolic preproteins from Hsp70/Hsp90 to the translocase of the outer membrane (TOM) complex. In humans, the cytosolic domain of Tom70 (*Hs*Tom70c) is entirely *α*-helical and comprises modular TPR motifs divided into an N-terminal chaperone-binding domain (NTD) and a C-terminal preprotein-binding domain (CTD). However, the mechanisms linking these functional regions remain poorly understood. Here, we present the 2.04 Å crystal structure of unliganded *Hs*Tom70c, revealing two distinct conformations – open and closed – within the asymmetric unit. These states are stabilized in part by interdomain crystal contacts and are supported in solution by hydrogen–deuterium exchange mass spectrometry (HDX-MS) and molecular dynamics (MD) simulations. Principal component and dynamical network analyses reveal a continuum of motion linking the NTD and CTD via key structural elements, notably residues in helices *α*7, *α*8, and *α*25. Engagement of the CTD by the viral protein Orf9b interrupts this network, stabilizing a partially-closed intermediate conformation and dampening dynamics at distal NTD sites. Collectively, our findings lay the groundwork for understanding Tom70 allostery and provide a framework for dissecting its mechanistic roles in chaperone engagement, mitochondrial import, and viral subversion.

## Introduction

Mitochondria utilize their unique dual-membrane architecture and a specialized proteome to support essential cellular functions, including respiration, signaling, and cofactor biosynthesis.^1^ Uniquely, roughly 99% of mitochondrial proteins are nuclear-encoded and synthesized in the cytosol as unstable precursors (preproteins), requiring chaperone-mediated stabilization for trafficking to the outer mitochondrial membrane (OMM) for subsequent import.^2^

In fungi and animals, preprotein import typically proceeds through the translocase of the outer membrane (TOM) and translocase of the inner membrane (TIM) complexes that span the outer and inner mitochondrial membranes (IMM), respectively.^3^ The membrane-embedded 7-subunit core TOM complex includes the preprotein receptor Tom22, and coordinates with the more transiently associated receptors Tom20 and Tom70 – each possessing a single-pass trans-membrane domain (TMD) and a cytosolic domain that facilitates substrate binding for import through the Tom40 pore-forming subunit.^4–7^ While substrate overlap is observed amongst the receptors, Tom20 and Tom22 canonically recognize N-terminal mitochondrial targeting sequences (MTS), while Tom70 directly interacts with chaperones and binds internal targeting signals (iMTS) present in the diverse array of mitochondrial carrier preproteins that must be imported.^8–11^

The cytosolic domain of Tom70 (Tom70c) functions as a dynamic adapter that recognizes preproteins bound to Hsp90 or Hsp70-family chaperones, and guides them to the TOM complex for mitochondrial import – a role reflected in its modular *α*-helical structure composed of 11 tetratricopeptide repeat (TPR) motifs.^12^ TPR1-TPR3 form the N-terminal domain (NTD) responsible for engagement of chaperone proteins,^13^ while TPR4-TPR11 comprise the C-terminal domain (CTD) that binds and stabilizes preprotein substrates.^14,15^ Structural studies of the cytosolic domain of yeast Tom71 (*Sc*Tom71c) – a paralog of *Sc*Tom70 – provide direct evidence of this modular function.^16,17^ These structures capture the NTD bound to C-terminal acidic chaperone fragments, with the CTD adopting an expanded conformation relative to apo-NTD *Sc*Tom71c. This shift was interpreted as a dynamic response to incoming preprotein cargo during handoff to the TOM complex, though the underlying mechanism remains incompletely understood.

Beyond its role in preprotein import, Tom70 also integrates mitochondrial function with cellular stress and immune responses, including mitochondrial morphology and involvement in endoplasmic reticulum-mitochondria contact sites, as well as the mitochondrial protein quality control and the mitochondrial antiviral signaling (MAVS) pathway in humans.^18–22^ Interest in Tom70 ‘s role in immune function surged during the SARS-CoV-2 pandemic, when the viral accessory protein Orf9b was shown to bind Tom70 ‘s preprotein-binding site, displacing Hsp90 and impairing the MAVS pathway.^23^ Orf9b is hypothesized to mimic endogenous MTS, adopting a unique *α*-helical conformation within Tom70 ‘s preprotein binding cleft, allosterically inhibiting its chaperone binding activity; however, the details of this apparent allostery remain unclear.^24–26^

Despite these findings, key questions about the regulation of Tom70 remain unexplored. While structural and biochemical experiments continue to indicate that allostery underpins Tom70 function in import, explanations for its underlying mechanism remain suggestive and ambiguous. In particular, the precise architecture and motions that couple NTD and CTD functions, giving rise to Tom70 ‘s allosteric behavior, are still largely speculative. Further, the absence of an unliganded *Hs*Tom70 structure has confounded efforts to address these questions, as all existing structures of *Hs*Tom70 are in complex with the viral inhibitor Orf9b. Thus, researchers have drawn comparisons to yeast homologs (*Sc*Tom70/71) to contextualize findings; however, these allusions are complicated by substantial conformational and subtle domain architectural differences between the two homologs, offering only a partial view of how Orf9b truly modulates *Hs*Tom70 structure and dynamics. Moreover, mechanistic parallels between Orf9b and endogenous presequence binding remain unclear without structural or dynamic data for unliganded *Hs*Tom70.

To address these questions, we characterized the dynamic landscape of the cytosolic domain of human Tom70 (*Hs*Tom70c) using X-ray crystallography, hydrogen–deuterium exchange mass spectrometry (HDX-MS), and molecular dynamics (MD) simulations. Crystallography reveals two distinct conformations within the asymmetric unit, representing “open” and “closed” conformations of *Hs*Tom70c that are potentially stabilized by contacts between adjacent molecules. HDX-MS and simulations further support the presence of these conformers in solution, enabling the construction of a putative allosteric network linking the NTD and CTD of *Hs*Tom70c. We applied these insights to the *Hs*Tom70c–Orf9b complex, using HDX-MS to contextualize existing structural data and assess the dynamic consequences of Orf9b binding. Moreover, our data suggest that Orf9b appears finely tuned to engage this dynamic network, stabilizing an intermediate *Hs*Tom70c conformation by remodeling the CTD and restricting Hsp90-interacting regions in the NTD to lock Tom70 in an inactive state. Our findings support a model in which intrinsic *Hs*Tom70c dynamics are essential for NTD-CTD coupling and, by extension, for Tom70 ‘s functional role in mitochondrial protein biogenesis, where binding at either the NTD or CTD alters the conformational landscape to ensure directionality in preprotein import.

## Results

### Crystal Structure and Solution Dynamics Reveal Conformational Plasticity in HsTom70c

To characterize the structure and dynamics of unliganded *Hs*Tom70c, we determined a 2.04 Å-resolution crystal structure of *Hs*Tom70c and performed HDX-MS analysis achieving 98% total sequence coverage **(Figure 1A–C, Supplementary Figure 1A, Supplementary Tables 1-2)**. *Hs*Tom70c comprises 25 *α*-helices, of which 22 assemble into 11 TPR motifs. Helices *α*1–7 constitute the NTD, which is connected to the CTD (helices *α*8–25) via the NTD-CTD linker. The electrostatic clasp that is essential for chaperone engagement resides within the NTD, while the CTD forms much of the preprotein binding cleft. Non-TPR helices *α*7, *α*8, and *α*25 serve as bridging elements that help organize the interface between the NTD and CTD and contribute to the architecture of the preprotein binding cleft.

**Figure 1.**
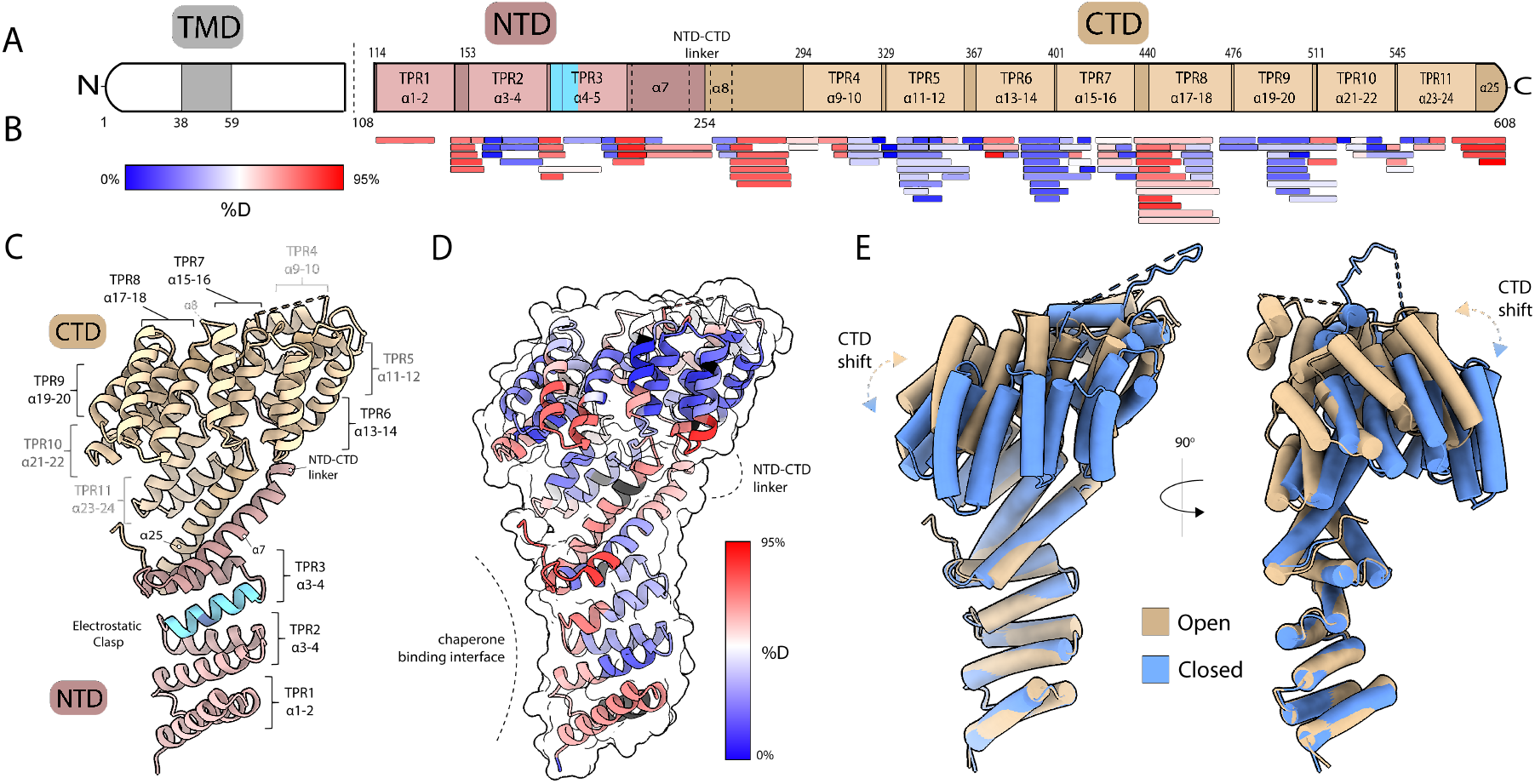
Structure and dynamics of the cytosolic domain of *Hs*Tom70. (**A)** Topology diagram of *Hs*Tom70 showing the transmembrane domain (TMD; grey), N-terminal domain (NTD; rose brown), and C-terminal domain (CTD; tan). Tetratricopeptide repeat (TPR) motifs and helices are also indicated. Dashed line at residue 108 indicates cytosolic domain of *Hs*Tom70 (residues 108-608; *Hs*Tom70c) used in this study. **(B)** HDX-MS peptide coverage map of *Hs*Tom70c aligned to the topology diagram. Peptides are colored by relative fractional deuterium uptake at the 2-minute exchange time point (0% uptake: blue; 66% uptake: red).(C) Ribbon representation of the 2.04 Å *Hs*Tom70c crystal structure (“open” conformation) annotated according to panel A. **(D)** HDX-MS relative fractional deuterium uptake heatmap at the 2-minute exchange time point overlaid on the open conformation ribbon structure. The chaperone-binding interface and NTD-CTD linker region are highlighted. **(E)** Structural superposition of open (tan) and closed (cornflower blue) *Hs*Tom70c conformations, aligned by the NTD. Conformational shifts in the C-terminal TPR motifs are emphasized.

HDX-MS provides complementary solution-phase insight into the conformational dynamics of *Hs*Tom70c **(Figure 1D)**. When normalized to back-exchange corrected fractional uptake, elevated deuterium incorporation is observed in the NTD. Specifically, in the electrostatic clasp region which includes the critical arginine 192 residue and corresponds to sites previously implicated in primary Hsp90 binding via HDX-MS.^27^ Increased exchange is also evident in *α*7, the NTD-CTD linker, and helix *α*25, indicating increased dynamics or solvent accessibility in these regions and supporting their potential as loci for allosteric NTD-CTD interdomain communication. Additionally, elevated uptake is detected in solvent-facing loops between TPR helices of the CTD, consistent with overall conformational plasticity in this region. Collectively, these results suggest *Hs*Tom70c samples a continuum of conformational states through coordinated NTD-CTD domain motions.

### HsTom70c Crystal Contacts Recapitulate Features of Chaperone-Mediated Allosteric Modulation

Strikingly, the asymmetric unit of our unliganded *Hs*Tom70c crystal structure contains two distinct conformations: a “closed” state, in which the preprotein-binding cavity is more occluded, and an “open” state with a more solvent-accessible binding cleft **(Figure 1E)**. Notably both conformers are in an elongated conformation, agreeing with previous solution-state insight into the domain organization of *Sc*Tom70.^28^ To investigate the origin of the distinct *Hs*Tom70c conformers observed within the asymmetric unit, we analyzed the crystal packing environment and intermolecular contacts **(Figure 2A, B)**. Notably, the open-state molecule engages the flexible loop region (residues 288–292) of a neighboring closed-state molecule via its electrostatic clasp. This interaction is mediated primarily by residues E288, L290, and E291 of the flexible loop, which contact Y186, K188, F125, and R192 of the neighboring open chain, respectively **(Figure 2C)**. Notable hydrogen bonding with the backbone atoms of the flexible loop residues is contributed by N122, Q157, and N158 of the neighboring open chain molecule.

**Figure 2.**
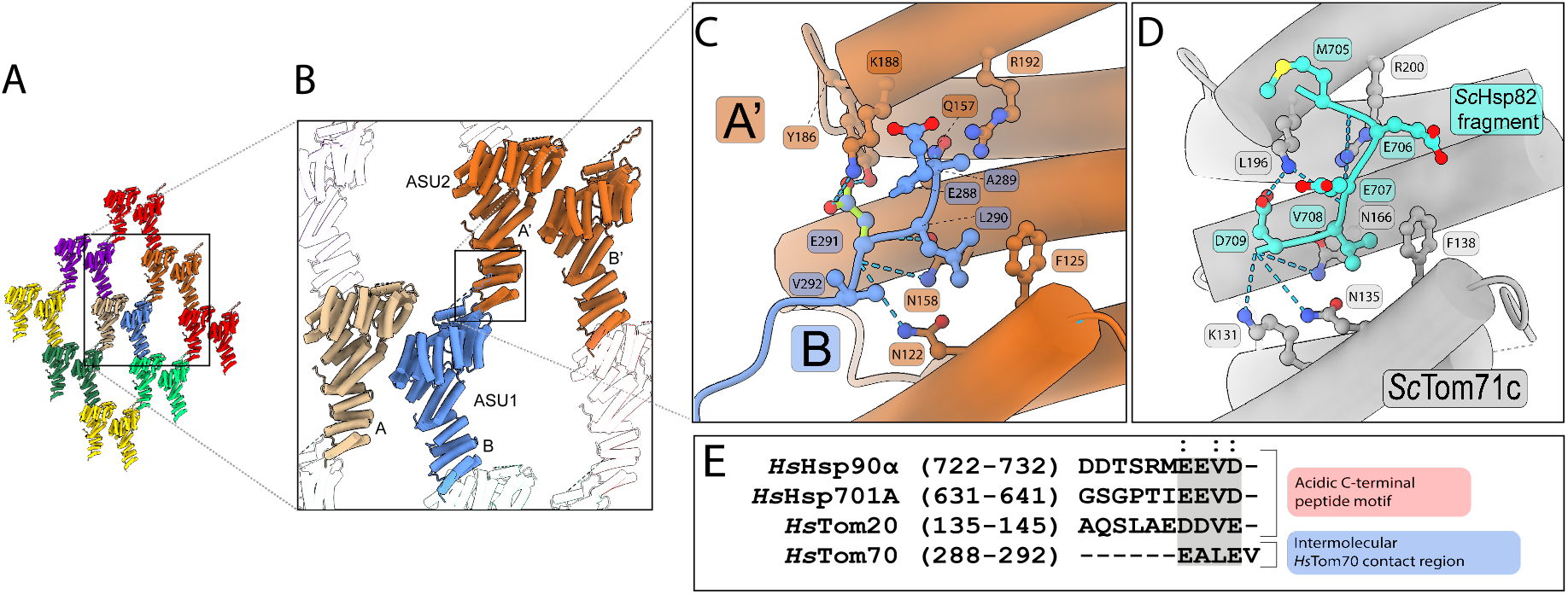
Crystallographic contacts between asymmetric units mimic acidic peptide engagement of the electrostatic clasp. **(A)** Arrangement of asymmetric units (ASUs) within the *Hs*Tom70c crystal lattice. **(B)** Interface between two neighboring ASUs, each containing one “open” (ASU1; chain A; tan) and one “closed” (ASU1; chain B; cornflower blue) conformation of *Hs*Tom70c. The disordered loop of the closed chain (chain B; cornflower blue) interacts with the electrostatic clasp of the adjacent open chain from a neighboring ASU (ASU2; chain A ‘; burnt orange). **(C)** Detailed molecular contacts mediating electrostatic clasp engagement. Residues 288-292 of the disordered loop from the closed chain (cornflower blue) form stabilizing interactions with clasp residues on the neighboring open chain (burnt orange). **(D)** Electrostatic clasp engagement of the acidic C-terminal tail of the yeast Hsp90 homolog *Sc*Hsp82 (cyan) with the yeast Tom70 homolog *Sc*Tom71c (light gray).

Although the biological relevance of this specific interaction remains uncertain, the binding mode closely resembles the highly conserved canonical acidic motifs of the EEVD sequence of Hsp70/Hsp90 and the DDVE motif of Tom20, and parallels interactions previously observed in *S. cerevisiae* Tom71 bound to C-terminal chaperone peptides **(Figure 2D, E)**.^9,16,17,29^ We hypothesize that clasp engagement by such motifs may modulate CTD dynamics, potentially priming the presequence-binding cavity for cargo interaction or release. Similar structural transitions have been reported in *Sc*Tom71 upon chaperone peptide binding, further supporting the functional relevance of these conformational states **(Supplementary Figure 2)**. Collectively, these observations underscore the intrinsic conformational flexibility of Tom70 homologs across distantly related species that could be important for preprotein binding, chaperone engagement, and directional preprotein import.

### Solution Dynamics Support an Open HsTom70c State Amid a Conformational Ensemble

To determine whether the conformations observed in our crystal structure represent accessible states within a broader conformational landscape and to contextualize our experimentally determined dynamics from our HDX-MS, we performed triplicate all-atom MD simulations initiated from both the “open” and “closed” conformations (see Methods). The resulting trajectories were overall similar in outcome, seemingly independent of the initial starting conformation; we therefore summarize the results concerning the open-state initiated simulations. The results of trajectories from both starting conditions can be found in the supplementary material **(Supplementary Figure 3, and Supplementary Movies 1-6)**.

To better understand our MD simulations with experimentally observed solution-phase HDX-MS dynamics, we applied HDX ensemble reweighting (HDXer), which adjusts the statistical weight of simulated conformers to better match experimental HDX-MS data.^30^ Using a maximum-entropy reweighting approach, our refined MD ensemble against the HDX-MS data resulted in substantially improved agreement with experimental uptake across most residues and time-points, indicating that the refined conformational landscape more accurately reflects *Hs*Tom70c ‘s apparent solution dynamics via HDX-MS **(Supplementary Figure 4 and 5)**.

To explore the structural characteristics of this optimized ensemble, we selected the top 1% of high-weight conformers and performed principal component analysis (PCA). Projecting these onto PC1 and PC2 revealed two primary populations identified by K-means clustering **(Figure 3A, Supplementary Figure 6)**. These clusters aligned with the experimentally derived “open” (tan) and “closed” (blue) conformers along PC1. Strikingly, the highest-weighted single frame (red X) most closely resembled the open conformation, suggesting this state offers the best overall agreement with solution-phase HDX-MS data **(Figure 3B)**.

**Figure 3.**
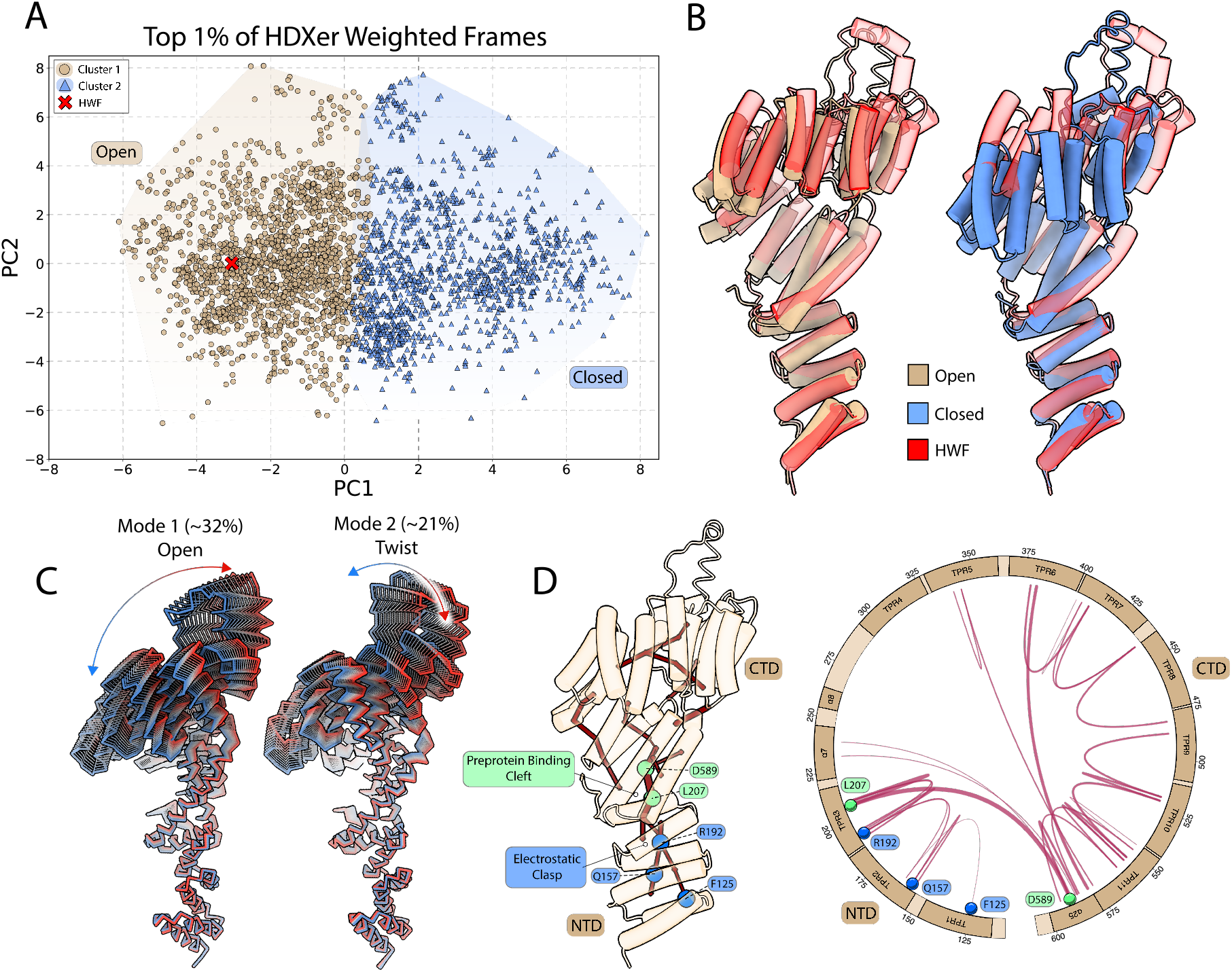
Molecular dynamics (MD) analysis of *Hs*Tom70c reveals coordinated motion and allosteric paths between the NTD and CTD. **(A)** Top 1% of weighted frames from HDXer analysis plotted in principal component (PC) space and mapped onto the experimentally determined open and closed *Hs*Tom70c structures. Two major clusters were identified along the PC mode 1 (PC1) axis, corresponding to open (dark green circles) and closed (light green triangles) conformations. Cluster centroids (white crosses) and the highest-weighted frame (HWF; yellow circle) are indicated. **(B)** Structural alignment of the HWF (yellow) with the open and closed *Hs*Tom70c structures. (C) PC modes 1 and 2 illustrate the dominant opening/closing and twisting motions of *Hs*Tom70c, respectively. The percentage of total observed motion captured by each PC is indicated. **(D)** Representative dynamical network mapped onto the open state crystal structure (cartoon, tan) and a corresponding circular topology plot of *Hs*Tom70c. Nodes and edges persistent in ≥ 75% of the trajectory are shown as red lines. Line thickness indicates more frequently observed interactions. Key residues involved in acidic peptide binding at the NTD (F125, Q157, R192) are highlighted as blue circles; L207 and D589 are highlighted as green circles.

### HsTom70c Samples a Continuum of Motion Linking Open and Closed Conformations

To characterize dominant motions within *Hs*Tom70c in solution, we conducted principal component analysis (PCA) on the concatenated MD trajectories. The first two principal components (PC1 and PC2) capture over 50% of the total conformational variance, accounting for ∼32% and ∼21%, respectively **(Figure 3C)**. Notably, these principal motions reflect two distinct global modes: an “opening” motion and a “twisting” motion of the CTD, which we refer to hereafter as the primary low-frequency collective movements inherent to *Hs*Tom70c. We interpret these results, together with our HDXer outcomes, as solution state evidence in support of the CTD dynamics inferred from our crystal structures.

### Dynamical Network Analysis Reveals the Internal Architecture Underlying Principal Motions and Allostery in HsTom70c

To further investigate the internal network dynamics underlying the principal motions identified by PCA in *Hs*Tom70c, we performed dynamical network analysis^31^ on our MD simulations to identify pathways of correlated motion across the protein **(Figure 3D)**. While the complete results are provided in the supplementary material **(Supplementary Figure 7)**, we distilled the analysis to highlight persistent nodes and edges – those present in at least 75% of the total trajectory, with edges weighted by their average betweenness. Several residues in the NTD, including F125, Q157, and R192 – previously implicated in binding acidic chaperone-derived peptides **(Figure 2C)** – connected distal regions of the NTD to the broader dynamic network. Notably, residues L207 of *α*6 and D589 of *α*25 at the base of the preprotein binding cleft at the NTD-CTD interface, exhibited strong correlated motion throughout the trajectory. This central pair connected with nodes and edges in the NTD, *α*7, and the CTD, suggesting their function as a principal conduit for interdomain communication. Together, these findings outline a putative allosteric pathway connecting correlated motions within the NTD to those in the CTD of *Hs*Tom70c. Further, our analysis suggests both open and closed conformations are a consequence of interdomain communication, supporting their functional importance.

### Global Remodeling of HsTom70c Dynamics Occurs Upon Orf9b Binding

Our characterization of the dynamic landscape of *Hs*Tom70c through both experimental and theoretical approaches afforded means to arrive at a putative allosteric network in the protein. We next examined how this allosteric architecture interacts with the SARS-CoV-2 accessory protein Orf9b, which suppresses the MAVS pathway by binding to *Hs*Tom70 and inhibiting its function.^23^ To complement existing structural insights,^24,25^ we expressed and purified the *Hs*Tom70c–full-length Orf9b complex **(Supplementary Figure 1B)** and used HDX-MS to assess changes in deuterium uptake in *Hs*Tom70c upon complexation **(Supplementary Table 2)**.

Compared to unliganded *Hs*Tom70c, the Orf9b-bound complex exhibited global dynamic remodeling, with up to ∼40% reduced deuterium uptake in the NTD **(Figure 4A)**. Strong protection was also observed in the preprotein-binding cavity where Orf9b resides *in crystallo*, aligning directly with contact sites in existing structures **(Figure 4B)**. Additional protection spans TPR3 and *α*7 of the NTD – extending toward a secondary Hsp90-binding site^26,27^ – and CTD elements TPR6, TPR7, TPR11, and *α*25. In contrast, increased uptake relative to apo-*Hs*Tom70c in *α*8 and the adjacent flexible loop, as well as TPRs 8-10, suggests localized CTD remodeling. While comparative HDX analyses of Orf9b were not performed, when internally compared and normalized against back-exchange corrected deuterium uptake across the protein, Orf9b-derived peptides show reduced exchange in the structurally informed binding interface with *Hs*Tom70c **(Supplementary Figure 8)**, consistent with successful complex formation.

**Figure 4.**
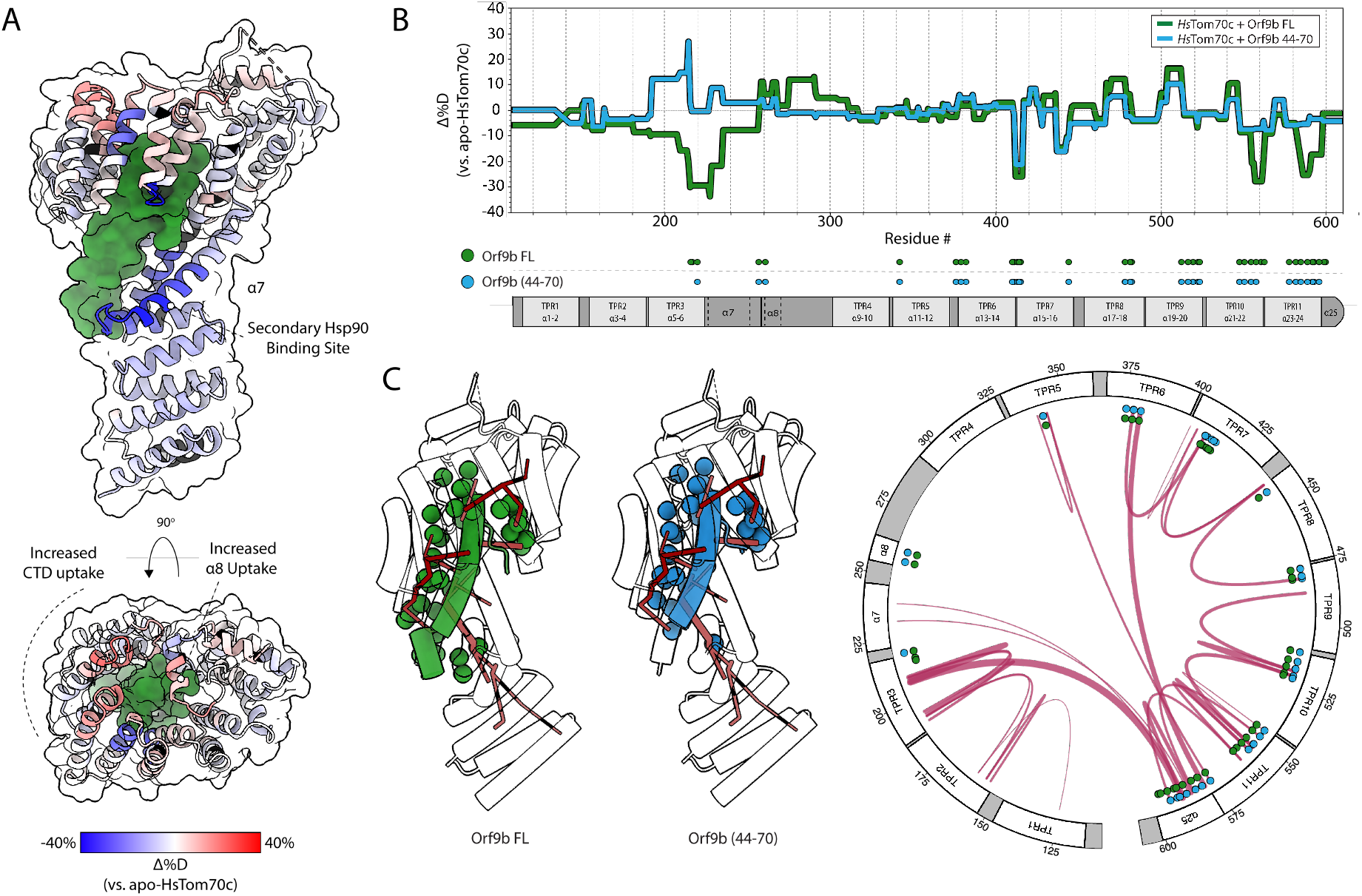
Orf9b tunes *Hs*Tom70c conformational dynamics via dual mechanisms of interaction. **(A)** Heatmap of relative fractional deuterium uptake differences between the *Hs*Tom70c-Orf9b complex and apo-*Hs*Tom70c at the 2-minute exchange time point (−40% uptake: blue; +40% uptake: red; no sequence coverage: black). Data are mapped onto a ribbon representation of *Hs*Tom70c (transparent surface) bound to Orf9b (forest green surface). **(B)** Butterfly difference plot showing fractional deuterium uptake relative to apo-*Hs*Tom70c for two complexes: *Hs*Tom70c + full-length Orf9b (Orf9b FL, green) and *Hs*Tom70c + Orf9b (residues 44-70; blue). The plot is aligned with a topology diagram of *Hs*Tom70c annotated with contact regions derived from crystal structures.**(C)** Crystal structure of *Hs*Tom70c bound to either full-length Orf9b (PDB: 7DHG, green) or Orf9b(44–70, blue), shown as transparent cartoons. Contact residues between *Hs*Tom70c and Orf9b are depicted as solid-colored spheres (FL, green; 44-70, blue) and mapped onto both the representative dynamic network (red lines) and a circularized topology plot.

### Differential Dynamics Induced by Orf9b Truncation Provide Insight Towards Allosteric Mechanism of Inhibition

To identify structural features critical for Orf9b-mediated modulation of *Hs*Tom70c, we performed HDX-MS using a previously characterized synthetic peptide comprising residues 44–70 of Orf9b,^25^ the minimally-observed Tom70 binding region, under saturating conditions **(Figure 4B, Supplementary Figure 9)**.^25^ When compared to exchange profiles of *Hs*Tom70c in complex with full-length Orf9b (Orf9b FL), most CTD regions exhibited similar dynamic trends. However, we noted reduced protection in TPR10 and TPR11, and markedly increased deuterium uptake in TPR3 and helix *α*7 of the NTD relative to *Hs*Tom70c-Orf9b FL. Notably, these dynamic alterations coincide with the loss of several interprotein contacts observed between *Hs*Tom70c and Orf9b FL *in crystallo*,^25^ suggesting that at least residues 71–78, absent in the truncated peptide, are essential for stabilizing the NTD.

To explore this further, we mapped the structurally informed contact sites associated with the resolved portion of Orf9b FL and the Orf9b (44-70) peptide onto our *Hs*Tom70c consensus dynamic network **(Figure 4C)**. Both Orf9b FL and Orf9b (44-70) engage all major CTD network nodes, thus suggesting a mechanism of inhibition where distal CTD interactions contribute NTD rigidification through long-range allosteric pathways. Further, we observe that minor alterations in protein-protein contact profiles between the fulllength and truncated forms of Orf9b lead to substantial differences in NTD dynamics, reinforcing a mechanism of inhibition dependent on Orf9b ‘s finely tuned allosteric modulation of *Hs*Tom70 via multifocal engagement.

### Orf9b Locks HsTom70c into A Partially Closed Intermediate Conformation

To further contextualize the structural changes induced by Orf9b binding, we compared the Orf9b-bound *Hs*Tom70c crystal structure^25^ with our open and closed conformations **(Figure 5A)**. Structural superposition via the NTD reveals minimal deviation; however, beginning at helices *α*7 and *α*8, significant divergence emerges and continues throughout the CTD **(Figure 5A)**. Among the various sites of perturbation, the *α*8 helix serves as a clear inflection point where structural differences between the open, closed, and Orf9b-bound states become pronounced **(Figure 5A-B)**. Notably, the *α*7 and *α*8 regions in the Orf9b-bound structure adopt conformations that are intermediate between those observed in the open and closed states, with a clear bias toward a partially closed configuration. These comparisons suggest that Orf9b binding stabilizes *Hs*Tom70c in a partially closed intermediate conformation, distinct from either crystallographic endpoint, consistent with a unique allosterically inhibited state.

**Figure 5.**
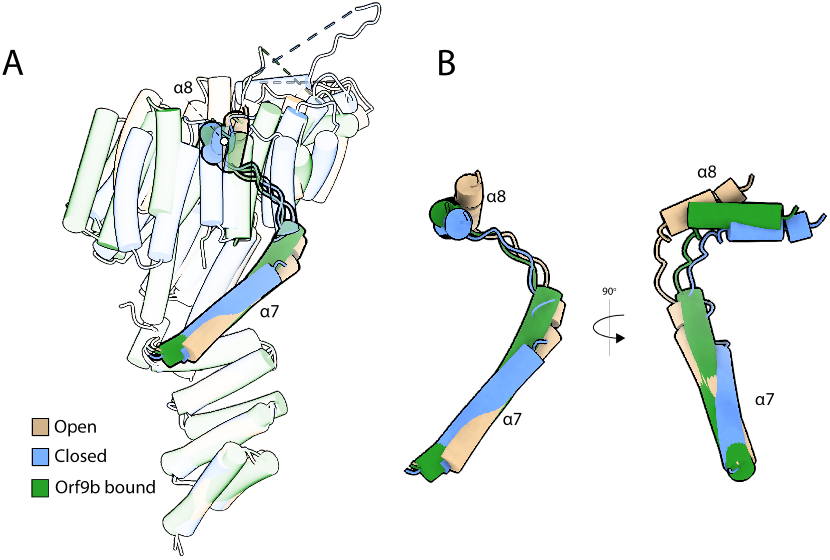
Orf9b locks *Hs*Tom70c into an intermediate conformation at a *α*7*α*8 hinge point. **(A)** Transparent overlay of the “open” state (tan), “closed” state (cornflower blue), and Orf9b-bound (forest green; PDB: 7DHG) structures of *Hs*Tom70c, highlighting the positions of the *α*7 and *α*8 helices in each structure of the alignment. **(B)** Inset showing the overlay of *α*7 and *α*8 from the “open”, “closed”, and Orf9b-bound *Hs*Tom70c structures.

## Discussion

We employed a combination of protein crystallography, HDX-MS, and all-atom molecular dynamics (MD) simulations to characterize the structure and dynamics of *Hs*Tom70c. Remarkably, our crystal structure captures two distinct conformations – designated “open” and “closed” – within the asymmetric unit **(Figure 1)**, distinguished by accessibility of the CTD preprotein-binding cleft and potentially stabilized *in crystallo* by intermolecular contacts at functionally relevant regions of the NTD **(Figure 2)**. HDX-MS data in combination with MD simulations describe two populations aligned with structural changes along the CTD and exhibit a clear bias toward an open-like conformation **(Figure 3)**. Further analyses support the presence of these dynamic oscillations of the CTD, revealing rapid opening and twisting motions that together account for over 50% of observed conformational variance and contribute to a dynamic communication network linking the NTD and CTD **(Figure 3)**.

We next applied these insights to investigate how *Hs*Tom70c is modulated by Orf9b, a viral inhibitor critical for SARS-CoV-2–mediated suppression of host immune signaling.^23–26^ HDX-MS analysis of the *Hs*Tom70c-Orf9b complex, alongside a synthetic Orf9b peptide, revealed extensive remodeling of both the NTD and CTD, largely dependent on residues 71-78 of Orf9b. Mapping Orf9b contact sites onto our computed dynamic network revealed broad engagement across key allosteric nodes, providing a mechanistic basis for its extensive influence on *Hs*Tom70c dynamics. Structural comparison of our open and closed conformers with the Orf9b-bound structure showed that most conformational changes localize to the CTD, with *α*7 and *α*8 marking a clear conformational inflection point. The Orf9b-bound conformation adopts a semi-closed intermediate state, distinct from either crystallographic endpoint.

Intramolecular contacts between asymmetric units in our crystal structure – particularly at the electrostatic clasp – offer further insight into how NTD conformational changes may influence CTD dynamics **(Figure 2)**. We speculate that interactions at this site helped stabilize the open conformation observed crystallographically. A similar phenomenon has been reported in *Sc*Tom71c structures, where CTD expansion was observed upon binding chaperone-derived fragments.^16,17^ Consistent with this, our HDX-MS analysis showed elevated deuterium uptake along a previously characterized chaperone-binding interface in the NTD,^27^ especially surrounding the electrostatic clasp. We interpret this increased exchange as evidence of conformational plasticity required for chaperone engagement and downstream communication with the CTD. Notably, residues such as R192 – critical for chaperone binding^9^ – also emerged as persistent nodes within our dynamic network, linking distal functional regions of the NTD and CTD **(Figures 3 and 4)**. These findings support a model in which acidic peptide binding in the NTD can allosterically modulate CTD structure and dynamics. The conservation of this dynamic coupling across species further highlights its functional significance in Tom70-mediated protein import.

To further understand how CTD perturbations influence distal regions of *Hs*Tom70c, we examined its interaction with Orf9b. HDX-MS revealed widespread deuterium protection across both domains, with localized increases in uptake suggesting rearrangement of C-terminal TPR motifs. Protection extended beyond direct contact sites into regions such as the electrostatic clasp and a secondary Hsp90-binding interface in the NTD.^26,27^ These data, considering prior isothermal titration calorimetric studies showing Orf9b inhibits Hsp90 acidic peptide binding,^25^ support a model in which Orf9b rigidifies flexible regions critical for chaperone recognition – functionally inactivating *Hs*Tom70c through long-range allosteric stabilization.

The portion of Orf9b resolved in available crystal and cryoEM structures represents only a segment of the full-length protomer.^24,25^ To evaluate whether unresolved regions contribute to *Hs*Tom70c dynamics, we performed HDX-MS using a previously characterized synthetic peptide spanning the preprotein-binding cleft.^25^ Although our experiments do not directly assess the function of the missing residues, we observed that the absence of residues 71-78 – present in the full-length protein but absent from the peptide – significantly impacted NTD stability. Specifically, HDX-MS revealed increased deuterium uptake across the NTD, consistent with a loss of stabilizing interactions. Structural comparisons indicate that the peptide recapitulates most full-length contacts, except for interactions in TPR3, TPR11, and *α*25. These findings suggest that Orf9b ‘s ability to allosterically stabilize *Hs*Tom70c depends, at least in part, on interactions mediated by residues 71–78, reinforcing their importance in modulating *Hs*Tom70c dynamics and inhibiting its function.

Finally, our dynamic network analysis reveals that Orf9b engages nearly all major nodes in *Hs*Tom70c ‘s internal communication pathway. Together, our data support a model in which Orf9b inhibits Tom70 function through a multifocal mechanism involving both direct occlusion of the preprotein-binding cleft and long-range remodeling of chaperone-binding interfaces. While further experiments are needed to dissect the individual contributions of these modes of inhibition, our findings provide a mechanistic framework and conceptual foundation for future investigations into Tom70 dynamics and viral interference.

We conclude by proposing a mechanistic model for substrate cycling by *Hs*Tom70 **(Figure 6)**, grounded in our integrated structural, biophysical, and computational data. In this model, the cytosolic domain of unliganded Tom70 extends from the outer mitochondrial membrane and dynamically samples conformational space. The electrostatic clasp of Tom70 engages preprotein-bound chaperones via the conserved acidic C-terminal EEVD motifs of cytosolic Hsp70/90 chaperones.^9^ This interaction initiates an allosteric relay that propagates conformational changes from the electrostatic clasp of Tom70 ‘s N-terminal domain (NTD) to the C-terminal domain (CTD), priming the substrate binding cleft to receive nascent preproteins from chaperones. Upon substrate handoff, preprotein engagement and the ongoing allosteric signal induce conformational rearrangements in the NTD and electrostatic clasp that lower chaperone affinity, promoting chaperone dissociation. We further propose that Tom70 ultimately transfers preproteins to Tom20, which engages the same electrostatic clasp region in the NTD via its acidic DDVE motif.^29^ This interaction is hypothesized to trigger a similar allosteric cascade, this time promoting substrate release for import via the TOM complex.

**Figure 6.**
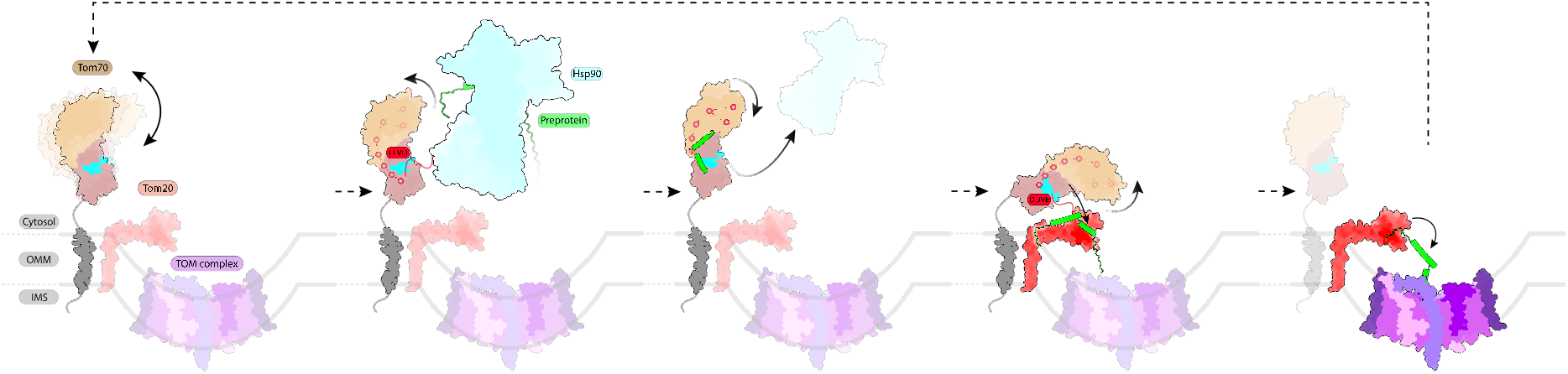
Simplified Model of Putative Tom70 Substrate Cycling. The cytosolic domain of apo Tom70 extends from the outer mitochondrial membrane and dynamically samples “open” and “closed” conformations. The electrostatic clasp of Tom70 engages preprotein-bound (sea green) chaperones (teal) via the conserved acidic C-terminal EEVD motifs of cytosolic Hsp70/90 chaperones. This interaction initiates an allosteric relay that propagates conformational changes from the electrostatic clasp of Tom70 ‘s N-terminal domain (NTD) to the C-terminal domain (CTD), priming the substrate binding cleft to receive nascent preproteins from chaperones. Upon substrate handoff, preprotein engagement and the ongoing allosteric signal induce conformational rearrangements in the NTD and electrostatic clasp that lower chaperone affinity, promoting chaperone dissociation. Preproteins are subsequently transferred to the TOM complex via Tom20 (red) via complexation of the electrostatic clasp of Tom70 through its own acidic DDVE motif, triggering Tom70 ‘s allosteric network to facilitate substrate release to Tom20 and subsequent import into the mitochondrion.

Our findings suggest that the conformational flexibility of *Hs*Tom70 is not merely an intrinsic property, but a functional requirement for coordinating chaperone engagement, preprotein capture, and substrate handoff. Our data suggest that CTD occupancy—natural or viral—can allosterically modulate NTD accessibility and support a model in which the directionality of substrate flux through Tom70 is governed by bidirectional allosteric coupling between the NTD and CTD.

## Methods

### Plasmid and Construct Design

The gene encoding full-length human Tom70 was codon-optimized for *E. coli* expression and synthesized by BioBasic (Canada). Two constructs encoding the Tom70 cytosolic domain (residues 108–608; *Hs*Tom70c) were prepared by subcloning into modified pET-24a vectors: one with an N-terminal His_6_–MBP tag followed by a 3C rhinovirus protease (3CRV) cleavage site, and another with a C-terminal 3CRV cleavage site followed by a tandem StrepII–His_6_ tag. The C-terminally tagged *Hs*Tom70c was used for crystallographic studies and Orf9b co-expression, while the N-terminally tagged *Hs*Tom70c was used for HDX-MS experiments of apo-*Hs*Tom70c or Orf9b (44-70) peptide incubated *Hs*Tom70c to minimize potential dynamic artifacts from C-terminal fusion in the absence of full-length Orf9b.

For the generation of the *Hs*Tom70c-Orf9b complexes, a co-expression construct was prepared as previously described. The full-length Orf9b gene was codon-optimized for *E. coli* expression, synthesized by GENEWIZ, and cloned into multiple cloning site 1 (MCS1) of a pET-DUET1 vector. The Tom70 cytosolic domain was inserted into MCS2 of the same vector and fused to a C-terminal 3CRV cleavage site followed by a tandem StrepII–His_6_ tag. All cloning approaches utilized Gibson assembly methods,^32^ primers were synthesized by Integrated DNA Technologies, and all constructs were sequence-confirmed by GENEWIZ.

### Recombinant Protein Expression and Purification

For crystallographic studies, the pET-24a vector encoding *Hs*Tom70c with a C-terminal 3C protease cleavage site and tandem StrepII-His_6_ tags was transformed into *E. coli* LOBSTR^33^ cells. Cultures were grown in terrific broth (TB) supplemented with 50 *μ*g/mL kanamycin at 37^*°*^C until an OD_600_ of ∼ 0.6–0.8. Protein expression was induced with 0.5 mM IPTG, followed by incubation at 18^*°*^C for 16–18 hours. Cells were harvested by centrifugation, resuspended in ice-cold Nickel Buffer A (20 mM Tris-HCl (pH 8.0), 500 mM NaCl, 10% glycerol, 25 mM imidazole-HCl (pH 8.0), 5 mM *β*-mercaptoethanol (BME), and supplemented with 1 mM MgCl_2_, 1 mM CaCl_2_, 1 mM PMSF, 5 mM benzamidine, 10 *μ*M leupeptin, 1 *μ*M pepstatin A, 2 *μ*g/mL aprotinin, and 0.1 mg/mL DNase I (Millipore Sigma). Cells were lysed by sonication on ice (Branson SFX550), and the lysate was clarified by centrifugation at 20,000 *×* g for 30 minutes at 4^*°*^C. All chromatography steps were carried out at 4^*°*^C, and SDS-PAGE was used to monitor elution fractions and assess protein purity.

The clarified lysate was loaded at 5 mL/min onto two tandem 5 mL HisTrap^*TM*^ HP columns (Cytiva) equilibrated in Nickel Buffer A. Columns were washed with 50 column volumes (CV) of buffer, and bound protein was eluted with 10 CV of Nickel Buffer B (20 mM Tris-HCl (pH 8.0), 150 mM NaCl, 500 mM imidazole-HCl (pH 8.0), 5 mM BME). Eluted fractions were pooled and passed through 10 mL of StrepTactin Superflow resin (IBA) equilibrated in Strep Buffer A (20 mM Tris-HCl (pH 8.0), 150 mM NaCl, 5 mM BME). After washing with 25 CV of Strep Buffer A, the protein was eluted with 5 CV of Strep Buffer B (Strep Buffer A plus 2.5 mM desthiobiotin). Eluted protein was pooled and incubated overnight at 4^*°*^C with 0.1 mg/mL His_6_-tagged 3CRV to remove the affinity tag. The reaction was adjusted to 500 mM NaCl and reapplied to the tandem HisTrap^*TM*^ columns equilibrated in Strep Buffer A to remove the His-tagged protease and cleaved tags. The flowthrough, containing the target protein with a C-terminal 3CRV protease tag scar, was collected. This flowthrough was diluted with 20 mM Tris-HCl (pH 8.0) to reduce NaCl to ∼ 75 mM, then applied to two tandem 5 mL HiTrap^*TM*^ Q HP anion exchange columns (Cytiva) equilibrated in Q Buffer A (20 mM Tris-HCl (pH 8.0), 75 mM NaCl, 1 mM dithiothreitol (DTT)). After washing with 5 CV of Q Buffer A, bound protein was eluted over a 20 CV linear gradient from 0% to 100% Q Buffer B (20 mM Tris-HCl (pH 8.0), 500 mM NaCl, 1 mM DTT). Selected fractions were pooled, concentrated, and loaded onto a HiLoad^*TM*^ 16/600 Superdex^*TM*^ 200 pg size-exclusion column (Cytiva) equilibrated in SEC buffer (20 mM Tris-HCl (pH 8.0), 150 mM NaCl, 0.5 mM tris(2-carboxyethyl)phosphine (TCEP). Homogeneous fractions were pooled and concentrated to ∼10 mg/mL. Samples were either used immediately or flash-frozen in liquid nitrogen with 10% glycerol and stored at −80^*°*^C.

For HDX-MS experiments, a pET-24a vector encoding *Hs*Tom70c with a N-terminal His_6_-MBP tag followed by a 3C protease cleavage site was used for the expression of apo-*Hs*Tom70c. The *Hs*Tom70c-Orf9b complex was expressed using a pET-DUET1 vector with Orf9b in MCS1 and *Hs*Tom70c in MCS2, bearing a C-terminal 3CRV cleavage site and tandem StrepII-His_6_ tag; expression and subsequent purification followed previously described methods with minor modifications.^25^ Each construct was transformed into *E. coli* LOBSTR^33^ cells. Cultures were grown in TB with 50 *μ*g/mL kanamycin for the cultures containing pET24a N-terminally tagged *Hs*Tom70c, and 100 *μ*g/mL ampicillin was used for cultures harboring the pET-DUET1 vector co-expressing Orf9b and C-terminally tagged *Hs*Tom70c. Cultures were grown at 37^*°*^C until an OD_600_ of ∼0.6–0.8. Expression was induced with 0.5 mM IPTG, and cells were incubated at 18^*°*^C for 16–18 hours. Cells were harvested by centrifugation, resuspended in ice-cold Nickel Buffer A (as described above) with protease inhibitors and DNase I, and lysed by sonication. Lysates were clarified at 20,000 × g for 30 minutes at 4^*°*^C. All purification steps were carried out at 4^*°*^C, with SDS-PAGE used to monitor protein purity. The clarified supernatant was loaded at 5 mL/min onto a 5 mL HisTrap^*TM*^ HP column (Cytiva) equilibrated in Nickel Buffer A. After washing with 50 CV, bound protein was eluted with 10 CV of Nickel Buffer B. Eluted fractions were pooled, treated with 0.1 mg/mL His_6_-3C protease, placed into dialysis tubing (3.5 kDa MWCO) (Thermo), and dialyzed overnight against 1 L of dialysis buffer (20 mM TrisHCl (pH 8.0), 150 mM NaCl, 5 mM BME. Following cleavage, the sample was adjusted to 500 mM NaCl and passed through two tandem 5 mL HisTrap^*TM*^ columns equilibrated in dialysis buffer to remove the protease and tag remnants. The flowthrough, containing protein harboring either an Nterminal 3CRV protease tag scar for apo-*Hs*Tom70c or a Cterminal 3CRV protease tag scar for *Hs*Tom70c in complex with Orf9b, was diluted to ∼ 75 mM NaCl and applied to a 5 mL HiTrap^*TM*^ Q HP columns equilibrated in Q Buffer A (20 mM Tris-HCl (pH 8.0), 75 mM NaCl, 1 mM DTT). After washing with 5 CV, the protein was eluted using a 20 CV gradient from 0% to 100% Q Buffer B (20 mM Tris-HCl (pH 8.0), 500 mM NaCl, 1 mM DTT). Final fractions were pooled, concentrated, and subjected to size-exclusion chromatography on a HiLoadTM 16/600 Superdex^*TM*^ 200 pg column equilibrated in HDX buffer (10 mM HEPES-NaOH (pH 8.0), 75 mM NaCl, 1 mM EDTA-NaOH (pH 8.0), 0.5 mM TCEP). Homogeneous fractions were concentrated to ∼10 mg/mL and either used fresh or flash-frozen in liquid nitrogen with 10% glycerol and stored at –80^*°*^C.

### Crystallization and Structure Determination of HsTom70c

For crystallization of *Hs*Tom70c, purified protein underwent lysine methylation by incubation with borane (50 mM final concentration) and formaldehyde (100 mM final concentration) for 1 hr at 4^*°*^C. The reaction was then quenched with 25 mM glycine (final concentration) for 30 min on ice. Methylated *Hs*Tom70c was then buffer-exchanged into crystallization buffer (25 mM Tris-HCl (pH 7.5), 200 mM NaCl, 5 mM MgCl_2_, and 1 mM TCEP) and concentrated to 16 mg/mL. Methylated *Hs*Tom70c was mixed in a 1:1 ratio with well solution containing 0.1M MES (pH 6.5), 0.1 M sodium acetate, and 30% PEG2000 MME in a sitting drop format. Crystals appeared after 24 hr and were harvested directly from the crystallization drop and frozen in liquid nitrogen.

Data were collected on beamline 24ID-C at the Advanced Photon Source at Argonne National Laboratory. Data were processed using the RAPD data processing pipeline (https://github.com/RAPD), which uses XDS^34^ for data indexing and reduction, AIMLESS^35^ for scaling, and TRUNCATE^36^ for conversion to structure factors. We determined the structure using molecular replacement in PHASER^37^ using an initial model generated by Alphafold 2.^38^ The model was manually rebuilt using COOT^39^ and refined in phenix.refine^40^ using positional, B-factor, and TLS refinement **(Supplementary Table 1)**.

### Hydrogen-Deuterium Exchange Mass Spectrometry (HDX-MS)

All samples were prepared for HDX-MS at an initial concentration of 10 *μ*M in HDX Buffer. For peptide mixing experiments, synthetic Orf9b (44-70) peptide (VWR) was resuspended in phosphate-buffered saline to a stock concentration of 2 mM. *Hs*Tom70c was maintained at 10 *μ*M and Orf9b (44-70) peptide was mixed to a working concentration of 90 *μ*M to ensure at least roughly ∼90% of *Hs*Tom70c was in complex with peptide at equilibrium after dilution during exchange.

HDX-MS was performed at the Biomolecular and Proteomics Mass Spectrometry Facility (BPMSF) of the University California, San Diego, using a Waters Synapt G2Si system with HDX technology (Waters Corporation) according to methods previously described.^41^ Briefly, deuterium exchange reactions were performed using a Leap HDX PAL autosampler (Leap Technologies, Carrboro, NC). D_2_O buffer was prepared by lyophilizing HDX buffer initially dissolved in ultrapure water and redissolving the powder in the same volume of 99.96% D_2_O (Cambridge Isotope Laboratories, Inc., Andover, MA) immediately before use. Deuterium exchange was measured in triplicate at each time point (0 min, 0.5 min, 1 min, 2 min, 5 min). For each deuteration time point, 4 *μ*L of protein was held at 25^*°*^C for 5 min before being mixed with 56 *μ*L of D_2_O buffer. The deuterium exchange was quenched for 1 min at 1^*°*^C by combining 50 *μ*L of the deuteration reaction with 50 *μ*L of 3M guanidine hydrochloride, final pH 2.66. The quenched sample (90 *μ*L) was then injected in a 100 *μ*L sample loop, followed by digestion on an in-line pepsin column (Immobilized Pepsin, Pierce) at 15^*°*^C. The resulting peptides were captured on a BEH C18 Vanguard precolumn, separated by analytical chromatography (Acquity UPLC BEH C18, 1.7 μm 1.0 *×* 50 mm, Waters Corporation) using a 7-85% acetonitrile gradient in 0.1% formic acid over 7.5 min, and electrosprayed into the Waters Synapt G2Si quadrupole time-of-flight mass spectrometer. The mass spectrometer was set to collect data in the Mobility, ESI+ mode; mass acquisition range of 200-2000 (m/z); scan time 0.4 s. Continuous lock mass correction was accomplished with infusion of leu-enkephalin (m/z = 556.277) every 30 s (mass accuracy of 1 ppm for calibration standard).

For peptide identification, the mass spectrometer was set to collect data in mobility-enhanced data-independent acquisition (MSE), mobility ESI+ mode instead. Peptide masses were identified from triplicate analyses and data were analyzed using the ProteinLynx global server (PLGS) version 3.0.3 (Waters Corporation). Peptide masses were identified using a minimum number of 250 ion counts for low energy peptides and 50 ion counts for their fragment ions. The following cutoffs were used to filter peptide sequence matches: minimum products per amino acid of 0.14, minimum score of 6.8, maximum MH+ error of 6 ppm, and a retention time RSD of 5%. The peptides identified in PLGS were then analyzed using DynamX 3.0 data analysis software (Waters Corporation). The relative deuterium uptake for each peptide was calculated by comparing the centroids of the mass envelopes of the deuterated samples with the undeuterated controls following previously published methods.^42^

For all HDX-MS data, at least 3 technical replicates were collected. Data are represented as mean values +/-STD of 3 technical replicates, though the LEAP robots high reproducibility may result in obscured error bars. The deuterium uptake was corrected for back-exchange using a global back exchange correction factor determined from the average percent exchange measured in disordered termini of various proteins.^43^ Deuterium uptake plots were generated in DECA,^44^ and the data are fitted with an exponential curve for ease of viewing. Peptides as shown in coverage map figures are the real coverage map data.

### System Preparation for Molecular Dynamics (MD) Simulations

The refined X-ray structures for the HsTom70c_*open*_ and HsTom70c_*closed*_ states were missing atoms in the *α*9-*α*10 loop (resides 272-296 for HsTom70c_*open*_ and 273-287 for HsTom70c_*closed*_). An ensemble of 20 models was generated using Alphafold3^45^ and these models were clustered using ChimeraX software (v1.9).^46^ The lowest energy model from the highest populated cluster for the *α*9-*α*10 loop, comprising residues 266-296, were then fit and transplanted to each starting structure prior to molecular dynamics (MD) simulations. Each model was solvated with explicit TIP3 water molecules and the number of Na^+^ and Cl^−^ ions were adjusted to neutralize the system charge and final ionic concentration set to 75mM to match HDX-MS experimental conditions. The final systems comprised a total of 264,008 and 264,125 atoms for the HsTom70c_*open*_ and HsTom70c_*closed*_, respectively, with Å^3^ orthorhombic periodic cells of 137 A for both systems.

All-atom MD simulations were performed using the CUDA memory-optimized version of NAMD 2.14^47^ and CHARMM36m force fields.^48^ Each simulation began with an energy minimization step to remove any steric clashes and ensure that the system was in a low-energy state. A minimization of 2000 steps with the following key settings: a cutoff of 12.0 Å for nonbonded interactions, a pair-list distance of 14.0 Å was performed and the Particle Mesh Ewald (PME) method for long-range electrostatics with a grid spacing of 1 Å was used.^49^ A switch distance of 10.0 Å was enabled and all atoms were wrapped to the primary simulation cell, including water molecules and ions, to maintain periodic boundary conditions. Following minimization, the system underwent a temperature annealing process by gradually increasing the temperature from 60 K to 300 K using Langevin dynamics with a damping coefficient of 1 ps^−1^ throughout the annealing phase. Each system was equilibrated at 300 K for a duration sufficient to reach a stable state. The equilibration phase used the same cutoff, pair-list distance, and PME settings as the previous steps to maintain consistency in the treatment of nonbonded interactions. Langevin dynamics continued to control the temperature, with a target pressure of 1.01325 bar maintained via a Langevin piston. The simulation time for this phase was 500,000 steps for both structures to ensure comprehensive equilibration. Harmonic constraints were applied during minimization, annealing, and equilibration to maintain the structural integrity of the starting complexes. Following minimization, heating, and equilibration, the systems were submitted for production MD simulations under NPT conditions. The MD production was carried out for 50,000,000 steps, equivalent to 100 ns, with coordinates saved every 1000 steps (2 ps). Randomized triplicates were conducted for each structure, yielding a combined total of 0.3 *μ*s (150,000 frames) per structure. The first 10 nanoseconds of each trajectory were excluded to ensure equilibrated data for all subsequent analyses **(Supplementary Figure 4)**.

### Essential Dynamic Analysis (EDA) of MD Trajectories

To identify the predominant modes of motion observed during molecular dynamics simulations, Essential dynamic analysis (EDA)^50^ was performed using the three 100-nanosecond concatenated trajectories from each system. The covariance matrix calculation used only protein C*α* atomic coordinates and was analyzed using the ProDy software package (v2.4.1).^51^ The covariance matrix of Cartesian coordinates was created from position fluctuations after subtracting the mean structure. Eigenvalues and eigenvectors were obtained by diagonalizing the covariance matrix. Top eigen-vectors were used to create dynamic animations and plots, showing correlated motions and major structural deformations.

### Dynamical Network Analysis (DNA)

DNA was performed following the methodology described by Luthey-Schulten and colleagues.^31^ Analyses were carried out using a Python-based framework (Python v3.9), employing the DNA package (v2.0) and MDAnalysis^52,53^ for trajectory processing. Atom ordering consistency and periodic boundary condition removal were achieved using the PrepareSystem function from the DNA package. For each system, the three 100-nanosecond trajectories were concatenated and divided into four consecutive 75-nanosecond windows. The correlation network for each window was derived from a subsample of ten uniformly spaced frames. Nodes were defined as C*α* atoms for amino acid residues and single pseudoatoms assigned to water molecules and ions where applicable. Residue pairs were connected by edges if the heavy atom distance between nodes remained within 4.5 Å for at least 75% of the simulation frames. Edge weights were calculated based on generalized correlation coefficients derived from mutual information estimates using Shannon entropy, with numerical performance optimized through NumPy and Numba.^54^

Network visualizations were generated in Visual Molecular Dynamics (VMD, v1.9.4-57a)^55^ using the custom NetworkView 2.0 graphical interface to highlight key residues and communities for clarity. The resulting visualizations were exported to UCSF ChimeraX (v1.9)^46^ for final stylization and figure preparation.

### HDX-MS-Guided Ensemble Refinement

To refine atomistic MD trajectories against our HDX-MS data, we used the HDXer open-source package (v1.2).^30^ This approach employs a maximum-entropy reweighting formalism, which minimally adjusts the weights of each MD frame to achieve optimal agreement between back-calculated and experimental deuterium uptake values. This process selects for a sub-ensemble of conformations that is most consistent with the experimental data while introducing the least possible perturbation to the original conformational distribution. The prior conformational ensemble was generated from all six independent, all-atom MD simulations (three closed-state, three open-state; 100 nanoseconds each). Solvent and ions were removed, and the coordinates were superposed on the reference crystal structure (C*α* atoms using MDTraj (v1.9.9).^56^ The dataset was downsampled to every fourth frame of the combined trajectory, yielding an input ensemble of 75,000 structures. The N-terminal methionine and the first amide of each peptide segment were excluded from the analysis to account for rapid back-exchange artifacts. The target dataset was derived from HDX-MS measurements performed at four exchange times (0.25, 0.5 1.0, and 2.0 mins). Back-exchange corrected deuterium uptake values were then converted to experimental protection factors under the assumption of EX2 exchange kinetics. For each frame in the input ensemble, theoretical protection factors were estimated using the empirical solvent-accessibility model implemented in HDXer. The reweighting was performed by using a hyper-regularization strength of *γ* equal to 2.0, a value determined from the “elbow” of an L-curve plot **(Supplementary Figure 5)**. This value optimally balances agreement with experimental data and preservation of ensemble diversity. The initial predictions from the unbiased simulation showed qualitative agreement but also notable deviations from the experimental measurements across the four labeling timepoints **(Supplementary Figure 6)**. As shown by the reweighted trace, the reweighting procedure yielded a better fit to the experimental deuteration fractions for most residues and time-points, confirming that the refined ensemble is a more accurate representation of the protein ‘s solution-state dynamics.

To characterize the structural ensemble that best fits the experimental HDX data, conformations corresponding to the top 1% of the final HDXer weights were selected for further analysis. This high-weight subset of frames was partitioned into two distinct conformational states using K-means clustering based on the C*α* atomic coordinates.

To visualize the relationship between these states, PCA was subsequently performed on the C*α* coordinates of the same set of frames. The structures were projected onto the first two principal components (PC1 and PC2), and the conformational space occupied by each cluster was delineated with a convex hull. For each cluster, a representative centroid structure was saved, defined as the frame with the minimum RMSD to the geometric center of its respective cluster. Additionally, the single conformation corresponding to the highest individual weight in the reweighted ensemble was identified and saved for further inspection.

## Inclusion & Ethics Statement

The research described here includes local researchers from the University of California, San Diego. The roles and responsibilities of this research were agreed upon by all included authors.

## Data Availability

The data supporting this study are available from the corresponding author upon request. All models have been deposited into the Protein Data Bank (PDB). The atomic coordinates of referenced structures are deposited in the PDB under the following codes:

PDB accession: 9PKQ

SBGrid Data Bank ID: 1186 | DOI: 10.15785/SBGRID/1186

All HDX-MS data referenced is available at massive.ucsd.edu under the following:

Data set: MSV000098542

Username: MSV000098542 reviewer

Password for web access: a

## Code Availability

The all-atom molecular dynamics simulations system files and parameter files are available at https://doi.org/10.5281/zenodo.14014935.

The custom code used in this study is available at https://doi.org/10.5281/zenodo.15882339 under the GPL-3.0 license. Detailed instructions for usage are provided in the repository ‘s README file. Additional scripts used for data preprocessing and figure generation are also included. For any further inquiries or collaborative use, please contact the corresponding author.

## Acknowledgements

We are grateful to the entirety of the Herzik lab for facilitating insightful discussions. We would like to thank the UCSD Physics Computing Facility for their computational support. Molecular graphics and analyses were performed with UCSF ChimeraX, developed by the Resource for Biocomputing, Visualization, and Informatics at the University of California, San Francisco (UCSF), with support from National Institutes of Health (NIH) grant R01-GM129325 and the Office of Cyber Infrastructure and Computational Biology, National Institute of Allergy and Infectious Diseases. This work was funded by the NIH grants R35-GM138206 (M.A.H), T32-GM008326 (M.J.B), and the UCSD Biomolecular and Proteomics Mass Spectrometry Facility through NIH shared instrumentation grant numbers S10 OD016234 (Synapt-HDX-MS) and S10 OD021724 (LUMOS Orbi-Trap). Additional support includes the Searle Scholars Program (M.A.H.), a George W. and Carol A. Lattimer UCSD Faculty Research Fellowship (M.A.H), and a Cottrell Scholars Award (M.A.H.). B.D.C is partly supported as a Goeddel Family Technology Sandbox Fellow. KDC acknowledges the National Institutes of Health (R35 GM144121).

## Author Contributions Statement

M.J.B. and M.A.H. conceived the work, oversaw data analysis and interpretation, and wrote the manuscript. M.J.B., E.A.K., K.D.C., and M.A.H. designed experiments. M.J.B. prepared samples. B.D.C., Y.Q., and K.D.C. conducted X-ray crystallography experiments, collected and analyzed data. M.J.B., S.S., and E.A.K. conducted HDX-MS experiments, collected and analyzed data. K.L.M. conducted and analyzed MD simulations. M.J.B., K.L.M., and M.A.H. prepared figures. All authors edited the manuscript.

