## Supplementary Materials for "An allosteric network governs Tom70 conformational dynamics to coordinate mitochondrial protein import"

**Table 1. Crystallographic Data Collection and Refinement for *Hs*Tom70c**

| | *Hs*Tom70c |
|---|---|
| **Data collection** | |
| Data collection date | 10/20/2022 |
| Beamline | APS 24ID-C |
| Wavelength (Å) | 0.97918 |
| Space group | $P2_1$ |
| Cell dimensions | |
| *a*, *b*, *c* (Å) | 65.45, 89.36, 105.78 |
| α, β, γ (°) | 90, 104.86, 90 |
| Resolution (Å)* | 102.24–2.00 (2.04–2.00) |
| $R_{merge}$* | 0.048 (1.006) |
| $I / \sigma I$ | 12.2 (0.8) |
| Completeness (%) | 97.6 (98.9) |
| Redundancy | 3.4 (3.5) |
| **Refinement** | |
| Resolution (Å) | 102.24 – 2.00 |
| No. reflections | 77515 |
| $R_{work}$ (%) | 0.2285 |
| $R_{free}$ (%) | 0.2605 |
| No. atoms | |
| Protein (non-H) | 7744 |
| Ligand/ion (non-H) | 0 |
| Water | 228 |
| *B*-factors | |
| Protein | 68.12 |
| Water | 58.52 |
| R.m.s. deviations | |
| Bond lengths (Å) | 0.005 |
| Bond angles (°) | 0.79 |
| Validation | |
| MolProbity score | |
| Clashscore | 6.20 |
| Poor rotamers (%) | 0.36 |
| Ramachandran plot | |
| Favored (%) | 98.34 |
| Allowed (%) | 1.66 |
| Disallowed (%) | 0 |
| Resolution (Å) | 102.24 – 2.00 |
| No. reflections | 77515 |
| PDB ID | 9PKQ |
| SBGrid Data Bank ID | 1186 |

*Values in parentheses are for highest-resolution shell.

**Table 2. Summary of HDX-MS Experimental Parameters and Dataset Quality Metrics**

| State File Dataset Name | Tom70 (108-608) | Tom70 (108-608) + Orf9b | Tom70 (108-608) + Orf9b (44-70) | Tom70 (108-608) + Orf9b |
|---|---|---|---|---|
| Protein Examined | *Hs*Tom70c | *Hs*Tom70c | *Hs*Tom70c | Orf9b |
| HDX Reaction Details | 10 mM HEPES-NaOH pH 8.0, 75 mM NaCl, 1 mM EDTA-NaOH, 0.5 mM TCEP, pD = 8.35, 25 °C | 10 mM HEPES-NaOH pH 8.0, 75 mM NaCl, 1 mM EDTA-NaOH, 0.5 mM TCEP, pD = 8.35, 25 °C | 10 mM HEPES-NaOH pH 8.0, 75 mM NaCl, 1 mM EDTA-NaOH, 0.5 mM TCEP, pD = 8.35, 25 °C | 10 mM HEPES-NaOH pH 8.0, 75 mM NaCl, 1 mM EDTA-NaOH, 0.5 mM TCEP, pD = 8.35, 25 °C |
| HDX Time Course (min) | 0, 0.25 , 0.50, 1.00, 2.00, 5.00 | 0, 0.25 , 0.50, 1.00, 2.00, 5.00 | 0, 0.25 , 0.50, 1.00, 2.00, 5.00 | 0, 0.25 , 0.50, 1.00, 2.00, 5.00 |
| HDX Control Samples | Maximally-labeled control | Maximally-labeled control | Maximally-labeled control | Maximally-labeled control |
| Back Exchange / Mean IQR | 28%/ 3% | 28%/ 3% | 28%/ 3% | 28%/ 3% |
| # of Peptides | 146 | 147 | 143 | 33 |
| Sequence Coverage | 98% | 97.60% | 98% | 100% |
| Average Peptide Length/ Redundancy | 14.42/4.29 | 14.37/4.24 | 14.30/4.16 | 17.31/4.82 |
| Replicates | 3 (technical) | 3 (technical) | 3 (technical) | 3 (technical) |
| Repeatability | ±0.15 Da (average standard deviation) | ±0.15 Da (average standard deviation) | ±0.13 Da (average standard deviation) | ±0.20 Da (average standard deviation) |
| Significance Criterion | ≥Δ0.5 Da | ≥Δ0.5 Da | ≥Δ0.5 Da | ≥Δ0.5 Da |

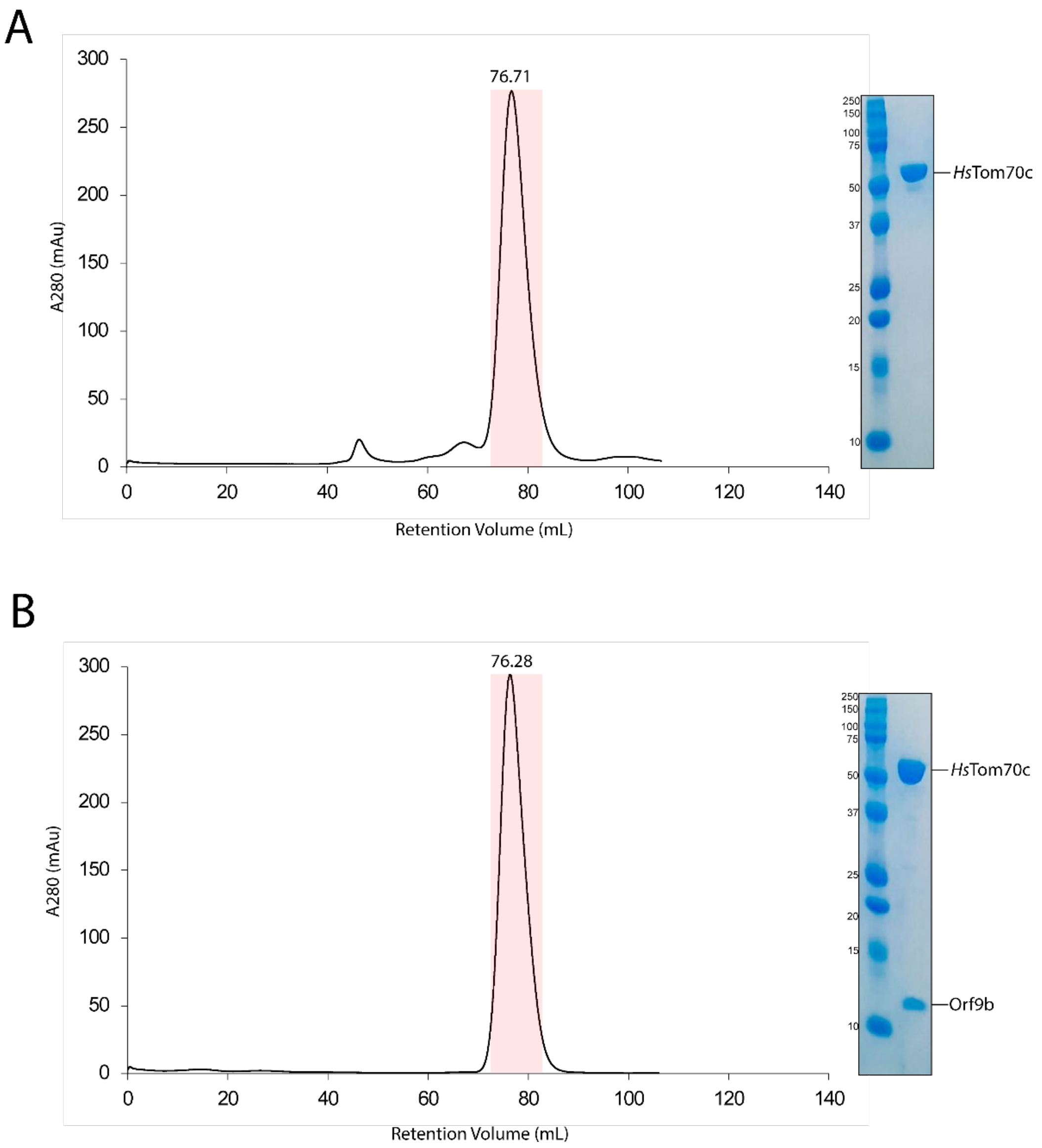

**Supplemental Figure 1 – Size-exclusion chromatographic analyses of *Hs*Tom70c and *Hs*Tom70c-full-length Orf9b purifications.**

Representative size-exclusion chromatogram of purified *Hs*Tom70c (A) and *Hs*Tom70c-full-length Orf9b (B) using a HiLoad 16/600 Superdex 200 pg are shown. Peak retention volume is annotated and fractions indicated by a red highlight were pooled. SDS-PAGE analysis of ~2 µg of estimated total protein.

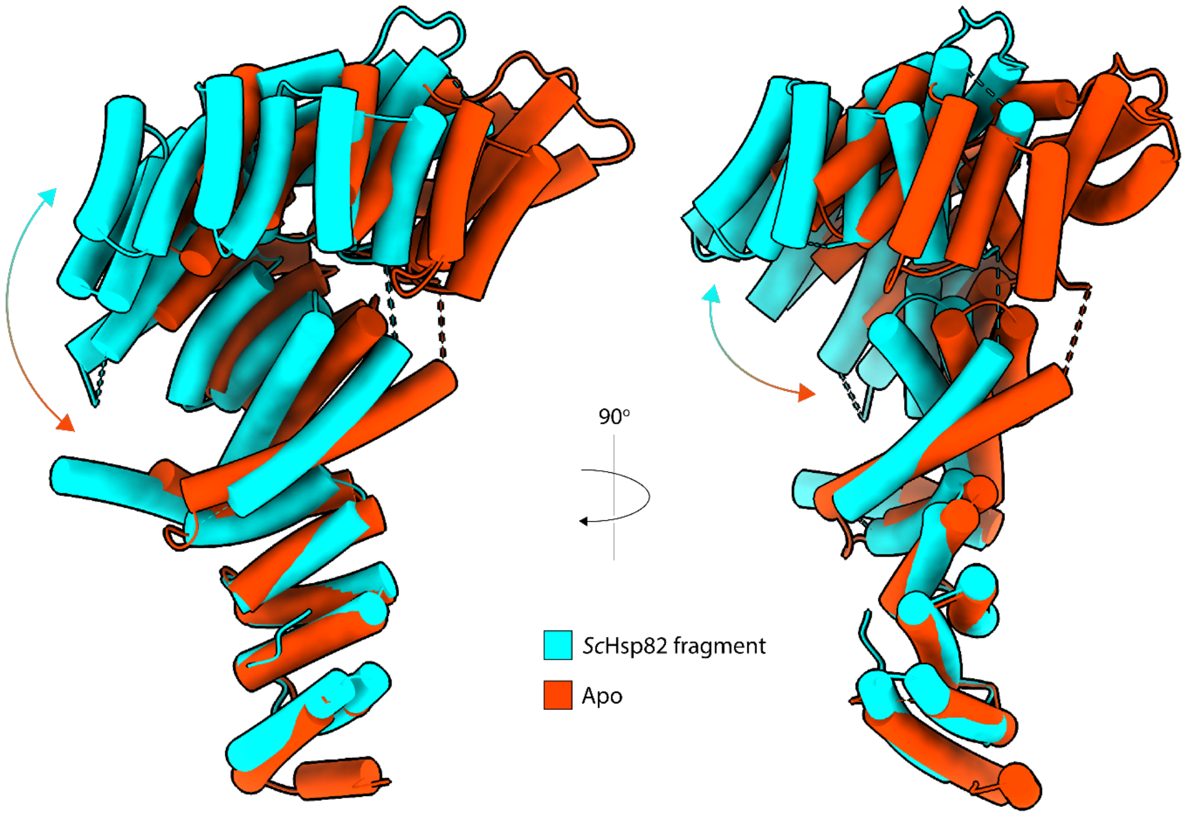

**Supplemental Figure 2 – CTD conformational change in *Sc*Tom71.**
Cartoon representation of the *ScTom71–ScHsp82* fragment crystal structure (PDB: 3FP2; cyan), aligned at the N-terminal domain (NTD) with the apo-*ScTom71* (PDB: 3FP3; orange).

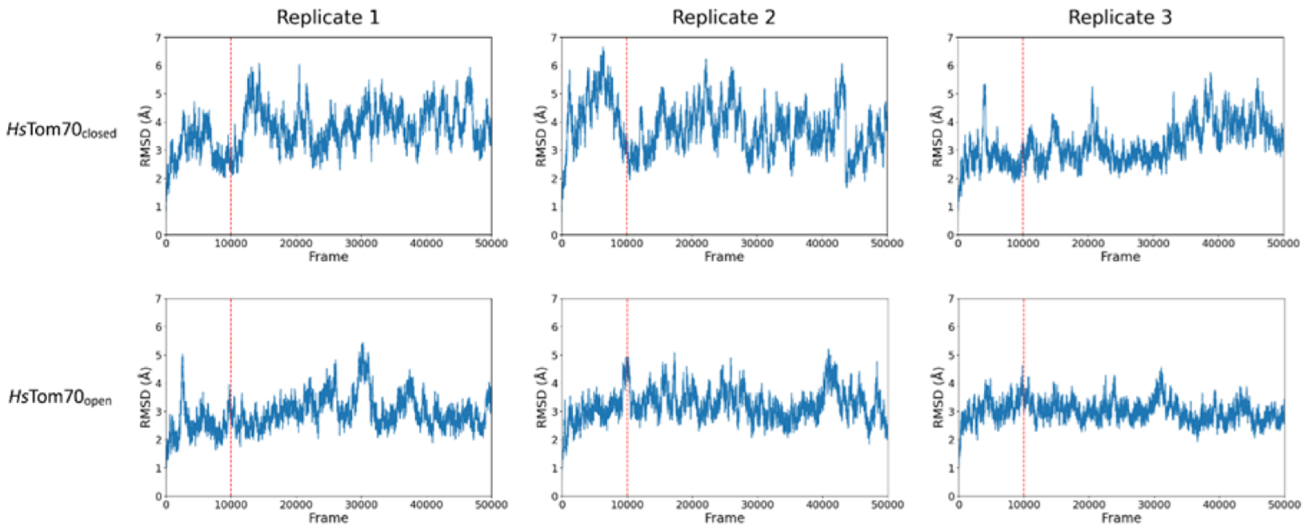

**Supplemental Figure 3 – RMSD analyses of MD trajectories.**
Plots from six independent 100 ns molecular dynamics (MD) simulations initialized from the *Hs*Tom70c closed (top row) and *Hs*Tom70c open (bottom row) conformation crystal structures. The vertical red dashed line indicates the cutoff for the 10 ns sections excluded from all analyses.

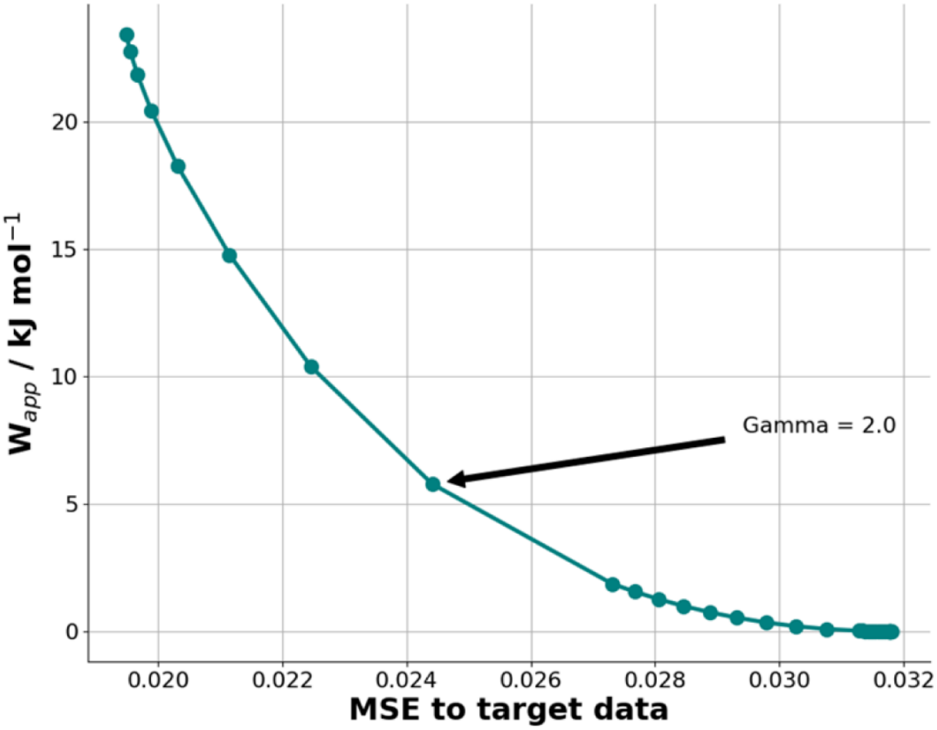

**Supplementary Figure 4 – *Hs*Tom70c HDXer decision plot.**
L-shaped curve plot illustrating the optimal γ (regularization parameter) selection for maximum-entropy reweighting of *Hs*Tom70 molecular dynamics ensembles to fit HDX-MS experimental data. The L-shaped curve shows an inflection point at γ value of 2.0, indicating an optimal value for reweighting that is reasonably balanced between fitting the experimental data and avoiding overfitting.

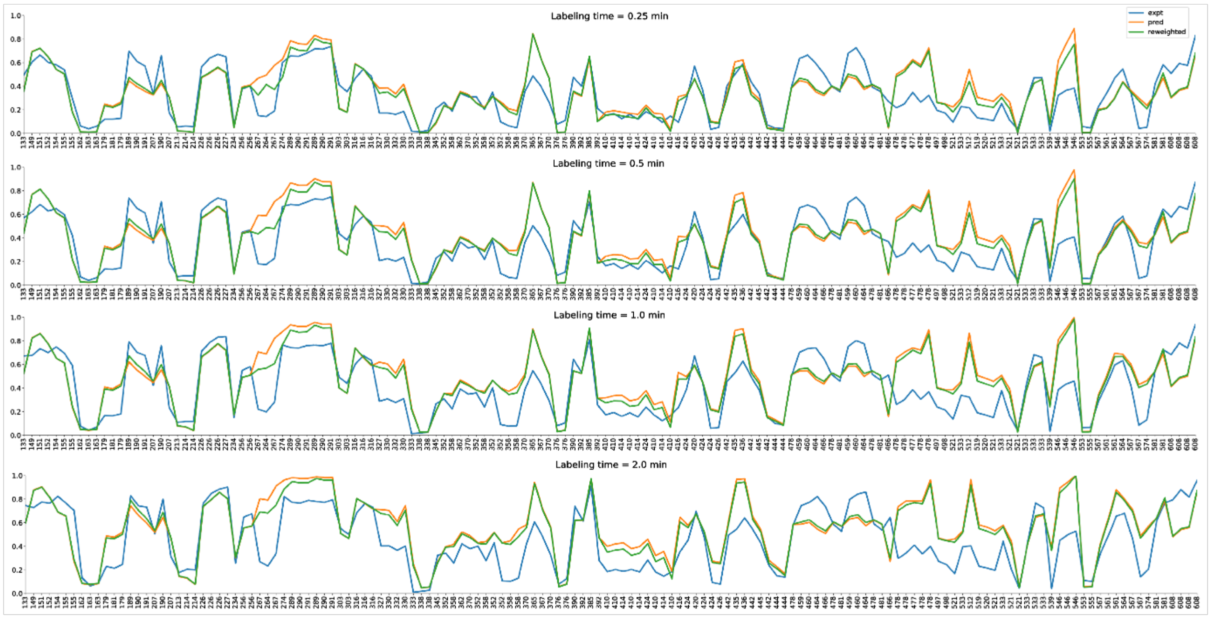

**Supplementary Figure 5 – *Hs*Tom70c HDXer reweighted deuterated fractions.**
Experimental deuteration values (blue) for *Hs*Tom70c residues are overlaid with predictions from the initial (orange) and reweighted (green) structural ensembles at various labeling times (0.25, 0.5, 1.0, and 2.0 min). The reweighted ensemble demonstrates improved agreement with experimental data across all time points compared to the initial ensemble.

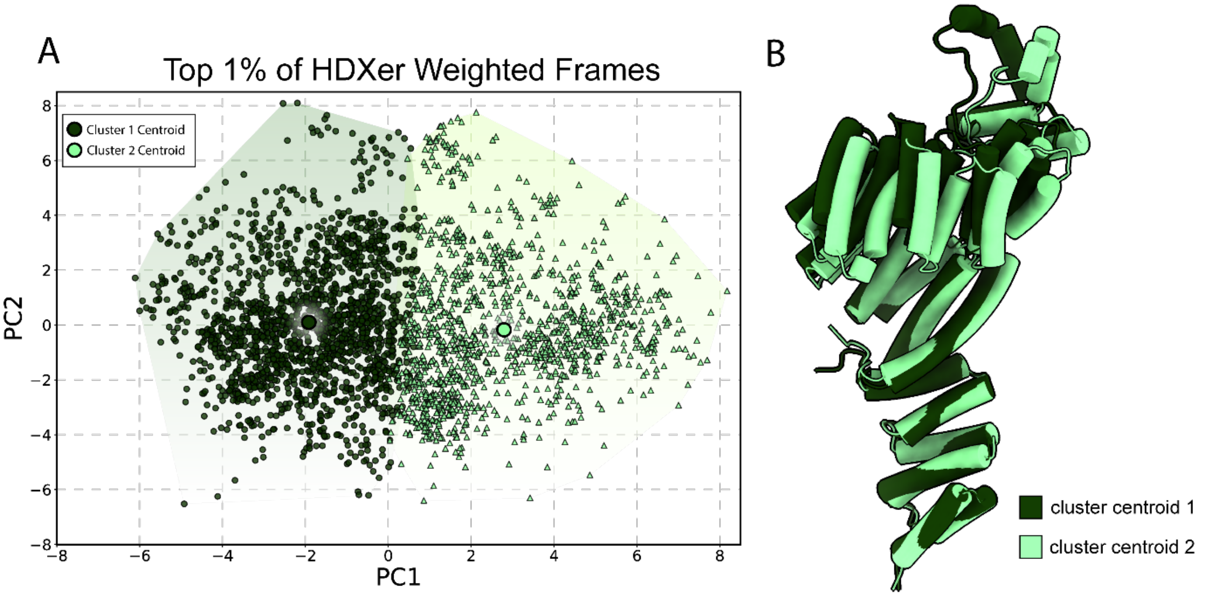

**Supplementary Figure 6 – Centroid structures of Distinct HDXer principal component Clusters.**

Top 1% of high-weighted HDXer frames projected into principal components 1 and 2. Light and dark green hulls indicate distinct clusters identified by K-means clustering, with centroids shown as large colored circles in PC space (A). Cartoon representations of centroid models from each cluster identified by K-means clustering of HDXer PCA results (B).

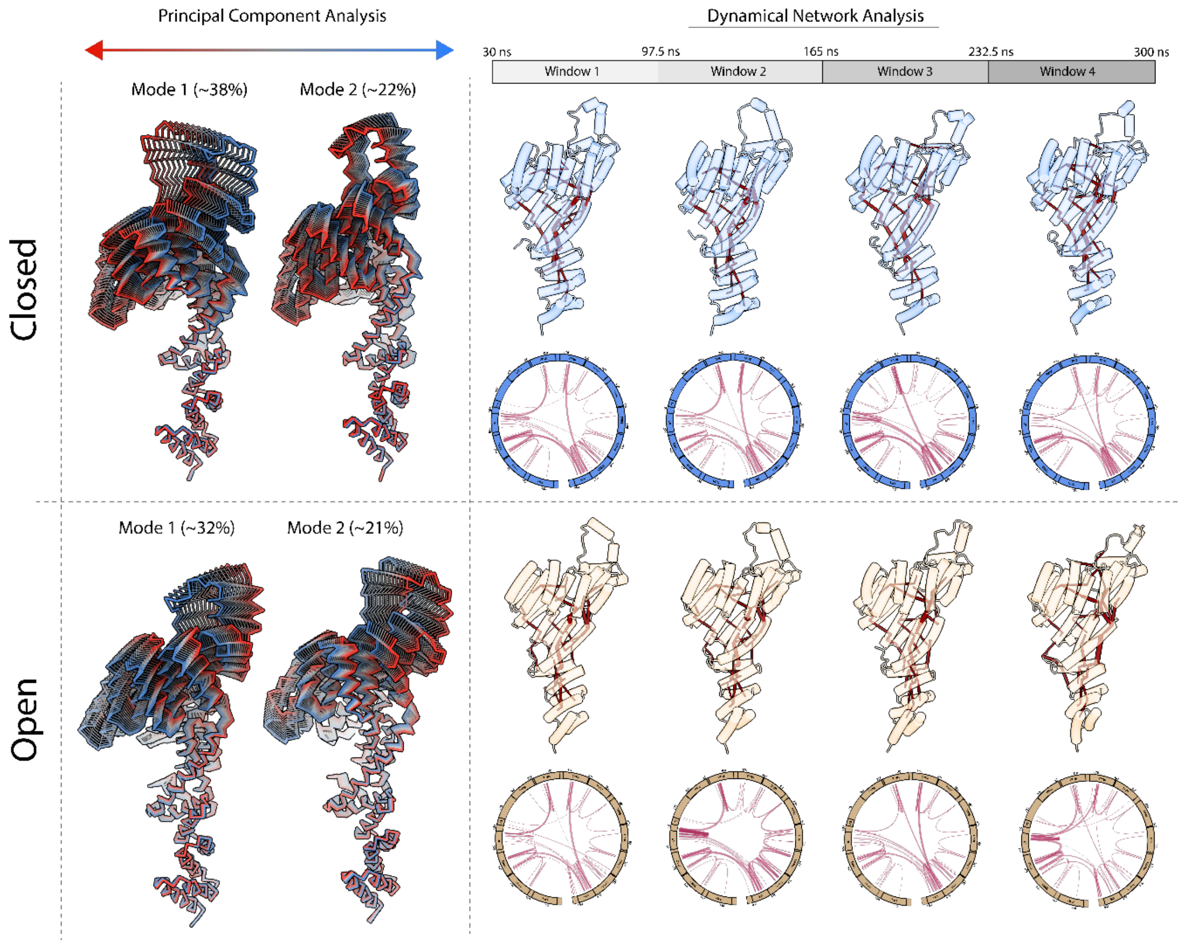

**Supplemental Figure 7 – Principal component and dynamical network analysis results initialized with open and closed starting models.**

*Left panels*: Closed (*top*) and open (*bottom*) conformations shown along principal component (PC) modes 1 and 2 from principal component analysis (PCA). Red denotes the starting structure; blue denotes the end point. *Right panels*: Representative structural snapshots from dynamical network analysis of the closed (*top*) and open (*bottom*) conformations. Snapshots are shown above corresponding 2D circular network plots of the concatenated trajectory. Nodes and edges are rendered as red lines with thickness corresponding to frequency of interactions.

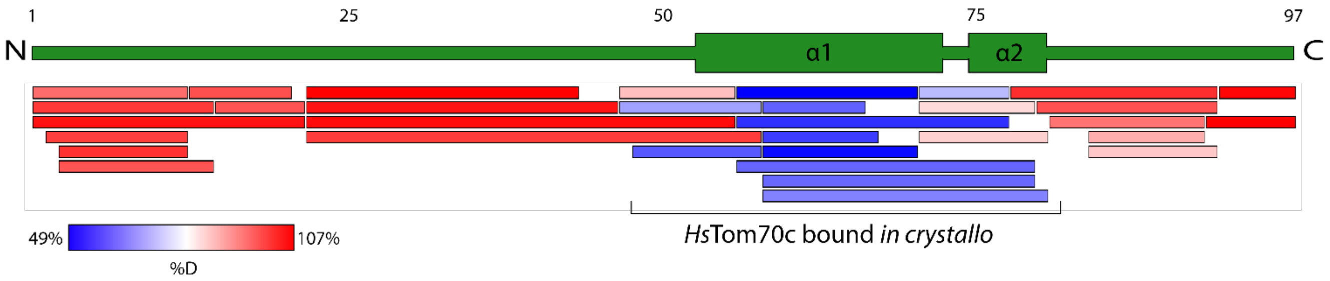

**Supplementary Figure 8 – Coverage map and relative fractional deuterium uptake of full-length Orf9b bound to *Hs*Tom70c.**

Topology cartoon (*top*; forest green) of full-length Orf9b bound to *Hs*Tom70c. The peptide coverage map and associated fractional uptake heatmap were generated using DynamX software with automatic scaling. Structured elements and regions that interact with *Hs*Tom70c *in crystallo* (PDB: 7DHG) are annotated.

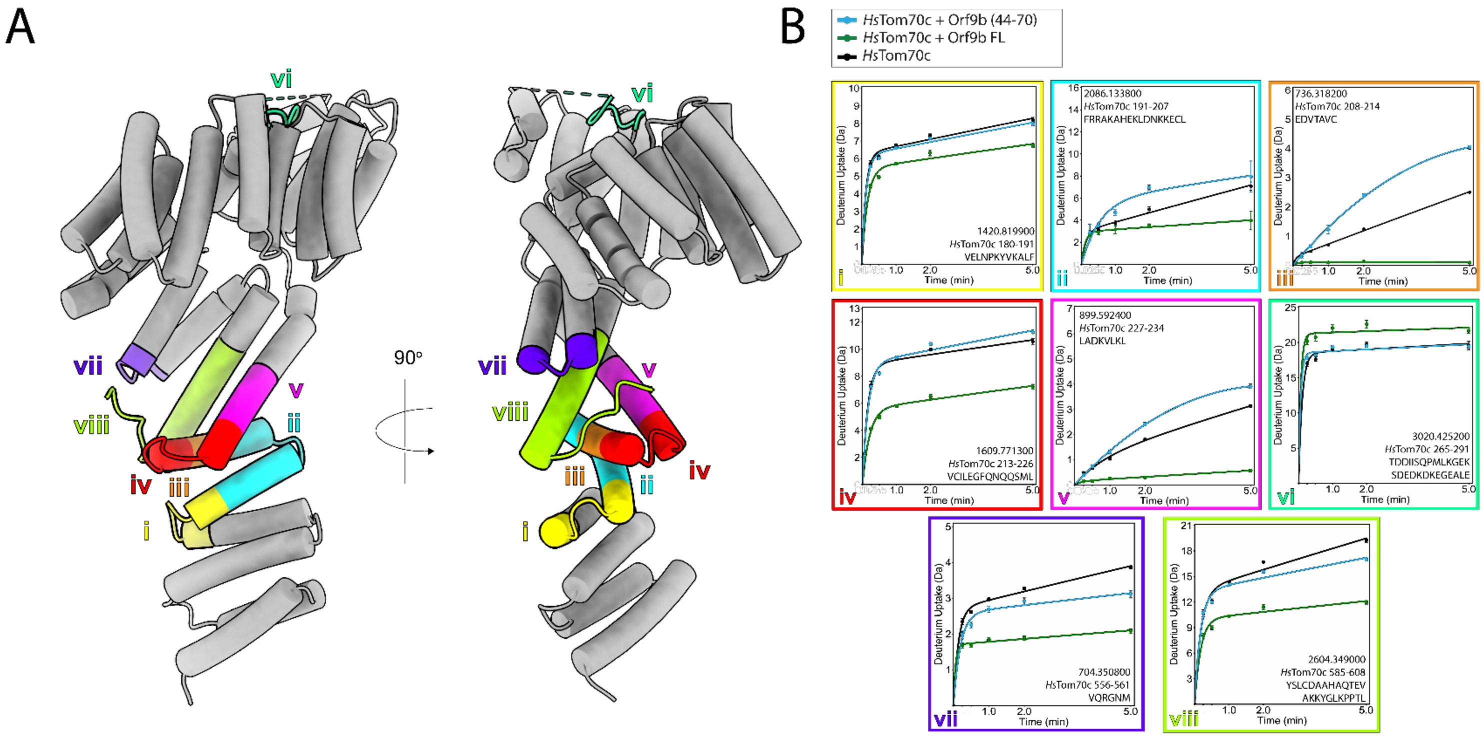

**Supplementary Figure 8 – Representative peptides highlighting distinct uptake profiles between full-length Orf9b and Orf9b (44–70).**

Cartoon representation (left) showing representative peptides mapped onto the structure, labeled by color and Roman numeral. Corresponding uptake plots (right) display deuterium uptake for each peptide using the same color and numeral scheme. Data points represent the mean of *n* = 3 replicates; error bars indicate standard deviation.
